# Neurostructural and molecular evidence of advanced brain aging in rheumatoid arthritis

**DOI:** 10.64898/2026.06.10.729182

**Authors:** Caroline Wasén, Malin C. Erlandsson, Karin Andersson, Kristian Stefanov, Amanda Gustafsson, Mina Hovland, Rille Pullerits, Sofia T. Silfverswärd, Kaj Blennow, Henrik Zetterberg, Neil Basu, Maria I. Bokarewa, Rolf A. Heckemann

**Affiliations:** Department of Rheumatology and Inflammation Research, Institute of Medicine, Sahlgrenska Academy, University of Gothenburg, Gothenburg, Sweden; Division of Rheumatology, Department of Medicine Solna, Karolinska Institutet, and Center for Molecular Medicine, Karolinska University Hospital, Stockholm, Sweden; Rheumatology Clinic, Sahlgrenska University Hospital, Gothenburg, Sweden; School of Infection & Immunity, University of Glasgow, Glasgow, United Kingdom; Department of Clinical Immunology and Transfusion Medicine, Sahlgrenska University Hospital, Gothenburg, Sweden; Department of Ophthalmology, Sahlgrenska University Hospital, Mölndal, Region Västra Götaland, Sweden; Department of Clinical Neuroscience, Institute of Neuroscience and Physiology, Sahlgrenska Academy, University of Gothenburg, Gothenburg, Sweden; Department of Psychiatry and Neurochemistry, Institute of Neuroscience and Physiology, Sahlgrenska Academy, University of Gothenburg, Mölndal, Sweden; Clinical Neurochemistry Laboratory, Sahlgrenska University Hospital, Mölndal, Sweden; Department of Neurodegenerative Disease, UCL Institute of Neurology, Queen Square, London, UK; UK Dementia Research Institute, UCL, London, UK; Hong Kong Center for Neurodegenerative Diseases, Clear Water Bay, Hong Kong, China; Wisconsin Alzheimer’s Disease Research Center, University of Wisconsin School of Medicine and Public Health, University of Wisconsin-Madison, Madison, WI, USA; Department of Medical Radiation Science, Institute of Clinical Sciences, Sahlgrenska Academy, University of Gothenburg, Gothenburg, Sweden; Department of Psychiatry and Neurochemistry, Institute of Neuroscience & Physiology, Sahlgrenska Academy, University of Gothenburg, Gothenburg, Sweden

**Keywords:** Rheumatoid arthritis, Brain atrophy, Neurodegeneration, Aging, Innate immunity

## Abstract

Chronic systemic inflammation has been implicated in age-related neurodegeneration, but whether rheumatoid arthritis (RA) is associated with accelerated brain aging remains unclear. We combined structural magnetic resonance imaging (MRI), circulating neurodegeneration biomarkers, and peripheral monocyte transcriptomics to investigate brain aging in RA across two independent cohorts. A brain-age prediction model trained in healthy controls from the IXI imaging dataset was applied to RA patient cohorts from Gothenburg (n = 71) and Glasgow (n = 50). RA was associated with significantly elevated corrected brain-age gap relative to healthy controls (+6.5 years, 95% CI 4.2–8.8 years, p = 1.2 × 10^−7^), with substantially stronger effects in patients ≥60 years. Older RA patients demonstrated a significant ventricular enlargement together with reduced frontal and parietal lobe volumes. Serum brain-derived tau and glial fibrillary acidic protein levels were elevated in RA. The increased brain-age gap was associated with altered myeloid transcriptional signatures. These findings demonstrate that RA is associated with age-related neurostructural alterations consistent with accelerated brain aging.

## Introduction

Aging leads to structural changes of the brain, including atrophy of cerebral white and grey matter resulting in enlargement of the brain ventricles ^1^, accelerating in the sixth decade of life ^2^. These processes occur in the normal aging brain but are often accelerated in patients with mild cognitive impairment and Alzheimer’s disease ^3^ and may be detectable before clinically evident cognitive decline and memory loss. Whether proper neuronal loss occurs in the healthy aging brain is a matter of controversy. Instead, cortical thinning and ventricular enlargement in the healthy brain are likely to be consequences of neuronal shrinkage ^4^, vascular changes ^5^, and possibly glial changes including astrocyte shrinkage. In Alzheimer’s disease, volume reduction of brain tissue is associated with amyloid β-induced neurodegeneration ^6^, leading to the release of brain-derived tau (BD-tau) and neurofilament protein (NFL), both of which can be measured in blood, serum or plasma to uncover cortical and subcortical neurodegeneration, respectively ^7,8^.

Systemic inflammation has been associated with age-related brain atrophy in cognitively unimpaired older elderly individuals ^9^, as well as with brain atrophy in patients with Alzheimer’s disease ^10^. Systemic inflammation induces neuroinflammation via the secretion of cytokines that induce microglia and astrocyte responses in the brain ^11^. In patients with mild cognitive impairment and Alzheimer’s disease, cortical volume reduction has been associated with microglial activation in affected regions ^12^. In neurodegenerative disease models, brain-resident macrophages adopt phenotypes referred to as disease-associated microglia (DAM) if their origin is yolk-sac derived microglia that populate the brain from birth, and disease inflammatory macrophages (DIM) if they stem from monocyte-derived macrophages that have migrated to the brain ^13^. Macrophages with the DIM signature have also been detected in single-cell RNA sequencing data sets of healthy aged mice, suggesting that monocytes may also migrate to the brain during normal aging ^13^. Macrophages of the DIM type have have been found in brain biopsies of patients with Alzheimer’s disease, but it is currently unknown whether they are present in the human aging brain in the absence of neuropathology ^13^.

Rheumatoid arthritis (RA) is a chronic systemic inflammatory disease that primarily affects the joints. Beyond its articular manifestations, structural brain abnormalities have been reported in patients with RA. We have previously found that smaller hippocampal volumes are associated with more severe RA ^14^, and that reduced premotor and supplementary motor area in the frontal cortex is associated with hand weakness^15^. Another study indicated that the hippocampus and orbitofrontal cortex are larger, and the thalamus is smaller in middle-aged female RA patients with mild cognitive impairment compared to age- and sex-matched healthy controls ^16^. A larger volume of the striatum in young adults was associated with a high genetic risk for RA, suggesting that pre-symptomatic processes of RA may induce brain volume abnormalities ^17^. Higher systemic inflammation in RA patients was associated with reduced grey matter volume in the inferior parietal lobule and posterior cingulate cortex ^18^, while fatigue correlated with larger putamen grey matter ^19^. More recently, large-scale neuroimaging analyses of UK Biobank data revealed increased white matter hyperintensity burden in RA, further supporting an association between chronic systemic inflammation and structural brain alterations ^20^. However, whether RA is associated with structural brain changes consistent with advanced brain aging remains unknown.

Here, we hypothesized that RA is associated with accelerated brain aging. To investigate this, we quantified regional brain volumes on magnetic resonance (MR) images using automated anatomical segmentation (MAPER ^21^), and applied these measures in a brain-age modelling framework. We further combined these analyses with blood-based biomarkers of neurodegeneration and neuroinflammation and transcriptomic profiling of peripheral CD14+ cells of these patients.

## Results

### Characteristics of the study cohorts

MRI analyses were performed in two independent cohorts of patients with RA, including a Gothenburg cohort (n = 71) and a Glasgow cohort (n = 50), together comprising 121 patients with established RA. Healthy reference data were obtained from the IXI dataset, which contains brain MR images of 581 healthy participants. IXI images were acquired at three London sites: Hammersmith Hospital (HH), Guy’s Hospital (Guys), and the Institute of Psychiatry (IOP). The Gothenburg cohort was older on average than the Glasgow cohort (mean age 59.2 versus 55.1 years), consisted exclusively of female participants, and displayed lower disease activity scores (mean 2.8 vs. 3.5 points, respectively). TNF inhibitor treatment was more common in the Gothenburg cohort, whereas disease duration was comparable between cohorts (**Table 1**). Despite these clinical differences, the two cohorts showed broadly comparable neurostructural findings across subsequent analyses.

**Table 1.** Socio-demographic and clinical data.

|  | RA<br>Gothenburg<br>n=71 | RA2<br>Glasgow<br>n=50 | Gothenburg vs Glasgow<br>Mann-Whitney/<br>Chi-square |
| --- | --- | --- | --- |
| Female | 100% | 76% | P<0.0001 |
| Age<br>(year) | Mean=59.2<br>SD=11.3<br>Range=23-76 | Mean=55.1<br>SD=11.4<br>Range=28-76 | P=0.023 |
| Disease<br>duration<br>(year) | Ave=13.5<br>SD=9.8<br>Range=0-45 | Mean =11.6<br>SD=8.8<br>Range=0.5-47.5 | ns |
| DAS28<br>(points) | Mean =2.8<br>SD=1.0<br>Range=1.1-5.8 | Mean =3.5<br>SD=1.2<br>Range=1.5-6.2 | P=0.0021 |
| Young-onset<br>RA | 44% | 30% | ns |
| RF positive | 75.4% | Unknown | NA |
| aCCP positive | 60.9% | Unknown | NA |
| MTX treated | 67.6% | 50.0% | ns |
| JAKi treated | 34.3% | Unknown | NA |
| TNFi treated | 38.0% | 14.0% | P=0.0038 |
| CTLA4 treated | 2.9% | Unknown | NA |
| IL6Ri treated | 4.3% | Unknown | NA |

RF, rheumatoid factor; aCCP, anti-cyclic citrullinated peptide; DAS28, disease activity score calculated by assessment of 28 joints; MTX, methotrexate; JAKi, Janus kinase inhibitors; TNFi, tumour necrosis factor inhibitors; CTLA4, Cytotoxic T-lymphocyte associated protein 4 fusion protein; IL6Ri, interleukin 6 receptor inhibitors.

### RI-based brain-age analysis reveals age-associated neurostructural alterations in RA

To investigate whether RA was associated with structural brain alterations consistent with accelerated aging, we trained the brain-age prediction model using MRI-derived regional brain volumes from healthy controls on the IXI imaging dataset. The model incorporated cortical, subcortical, posterior fossa, and ventricular regions and demonstrated moderate prediction accuracy in healthy controls (R^2^ = 0.53, r = 0.73; **Fig. S1A**). As expected for brain-age models, raw predictions displayed age-dependent bias, with overestimation of younger individuals and underestimation of older individuals. This bias towards the mean was effectively eliminated using spline-based correction, resulting in no residual association between corrected brain-age gap and chronological age in IXI controls (**Fig. S1B**). Leave-one-site-out validation across the control sites (HH, Guys, and IOP) demonstrated generally stable performance across scanners, although some site-dependent variation remained after the correction (**Fig. S1C**).

Application of the trained model to the combined RA cohort revealed a significantly elevated corrected brain-age gap relative to the healthy reference group (mean brain-age gap = +6.5 years, 95% CI 4.2–8.8 years, one-sample t-test p = 1.2 × 10^−7^). Stratification by age demonstrated that this effect was substantially more pronounced in older individuals (**Fig. 1B**). Patients younger than 60 years displayed a modest increase in corrected brain-age gap (+2.0 years, Wilcoxon p = 0.028), whereas patients older than 60 years showed a markedly elevated brain-age gap (+8.1 years, Wilcoxon p = 1.2 × 10^−7^).

**Figure 1.**
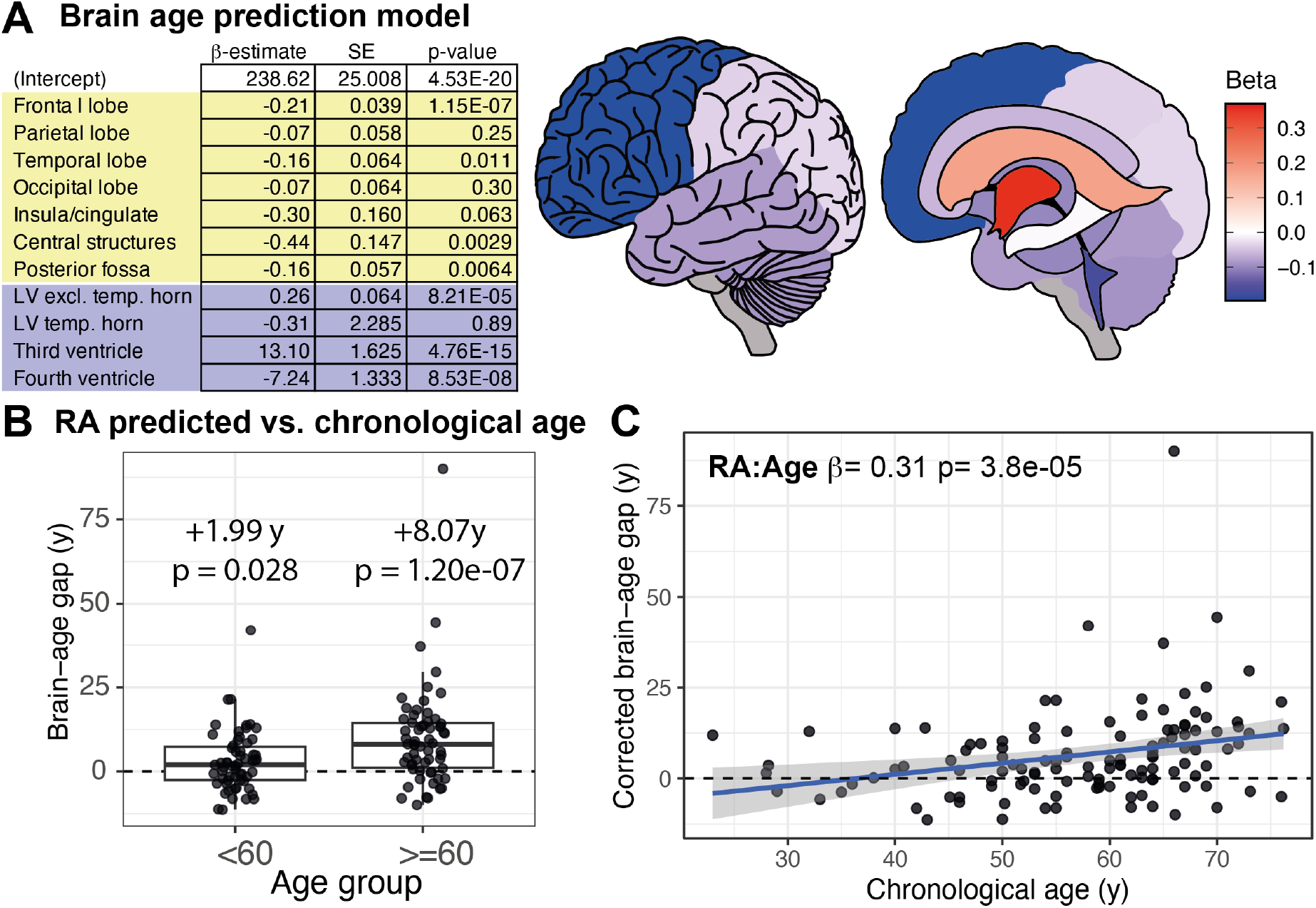
Brain-age prediction model and age-associated brain-age gap in rheumatoid arthritis (RA). **A)** Linear regression model trained in healthy controls from the IXI dataset (n = 566; HH, Guys, and IOP sites pooled) using intracranial volume-normalized MRI-derived brain region volumes to predict chronological age. The table shows standardized β-estimates, standard errors (SE), and p-values from the scaled regression model. The brain illustrations indicate associations between brain region volumes and age (scaled beta estimates from the regression model). **B)** Corrected brain-age gap in RA patients from the Gothenburg (n = 71) and Glasgow (n = 50) cohorts stratified by age (<60 years and ≥60 years). Corrected brain-age gap reflects the difference between predicted brain age and the brain age expected for an individual’s chronological age based on IXI healthy controls. Positive values indicate an older-appearing brain than expected for chronological age. P-values indicate deviation from zero corrected brain-age gap using two-sided Wilcoxon signed-rank tests. **C)** Association between chronological age and corrected brain-age gap in RA patients estimated using a linear regression model with corrected brain-age gap as the dependent variable and chronological age as the independent variable. Shaded area represents the 95% confidence interval of the linear regression model.

Corrected brain-age gap increased significantly with chronological age in RA patients (β = 0.31 years/year, p = 0.0022; **Fig. 1C**), whereas no corresponding age association was present in IXI healthy controls following bias correction. A formal diagnosis-by-age interaction test supported this difference between groups (p = 3.8 × 10^−5^), indicating that the age-related increase in corrected brain-age gap was specific to RA. A similar age-associated pattern was observed across both the Gothenburg and Glasgow cohorts despite differences in clinical characteristics (**Fig. S1D**).

Exploratory analysis was performed to assess associations between clinical and treatment variables and corrected brain-age gap within the combined RA cohort (**Fig. S2**). In age-stratified regression analyses, age at RA onset above 60 years was associated with increased brain-age gaps, whereas biological DMARD treatment was associated with diminution of the brain-age gap (**Fig. S2A**). Additionally, multivariable treatment modelling identified an interaction between methotrexate, TNF inhibitor (TNFi) treatment, and age group (**Fig. S2B**). Estimated marginal means from the regression model suggested that combined methotrexate and TNFi treatment was associated with smaller predicted brain-age gap in older patients compared with other treatment groups (**Fig. S2C**).

Further exploratory analyses were performed to assess whether cardiovascular risk factors contributed to the increased brain-age gap observed in RA (**Fig. S3A**). After age correction, overall Framingham cardiovascular risk scores showed no strong association with corrected brain-age gap within the RA cohorts. Hypertension status was associated with increased brain-age gap (**Fig. S3B**); however, elevated brain-age gap remained present in RA patients without hypertension compared with healthy controls (**Fig. S3C**).

### RA is associated with reduced frontoparietal brain volumes and enlarged ventricles

To identify the neuroanatomical alterations that contribute to the elevated brain-age gaps observed in RA, we next analyzed regional brain volumes in two independent RA cohorts (Gothenburg, n = 71; Glasgow, n = 50) and compared them with those of age- and sex-matched healthy controls from the multi-site IXI dataset (HH, Guys, and IOP). Given the residual site-dependent variation observed in the brain-age validation analyses (**Fig. S1C**), we explicitly quantified inter-site MRI variability within the IXI dataset and used this as a reference framework for interpreting disease-associated structural effects. For each brain region, we calculated the absolute effect size (Hedges’ g) for differences between RA patients and pooled IXI controls. Comparing this with the maximum pairwise inter-site effect size observed across IXI scanner sites enabled estimation of a signal-to-noise ratio.

In participants older than 60 years, RA-associated regional brain volume differences frequently exceeded inter-site MRI variability (**Fig. 2A**). Ventricular regions, particularly the lateral ventricles excluding the temporal horns, showed effect sizes larger than scanner/site variability in both cohorts, indicating robust ventricular enlargement in older RA patients. Among cortical regions, the frontal and parietal lobes demonstrated the most consistent reductions across cohorts, with disease-associated effect sizes exceeding inter-site variability. In contrast, temporal and occipital regions displayed smaller and less consistent differences. Corresponding analyses of patients younger than 60 years revealed a substantially weaker brain regional effect, with most brain regions showing effect sizes comparable to scanner/site variability (**Fig. S4**).

**Figure 2.**
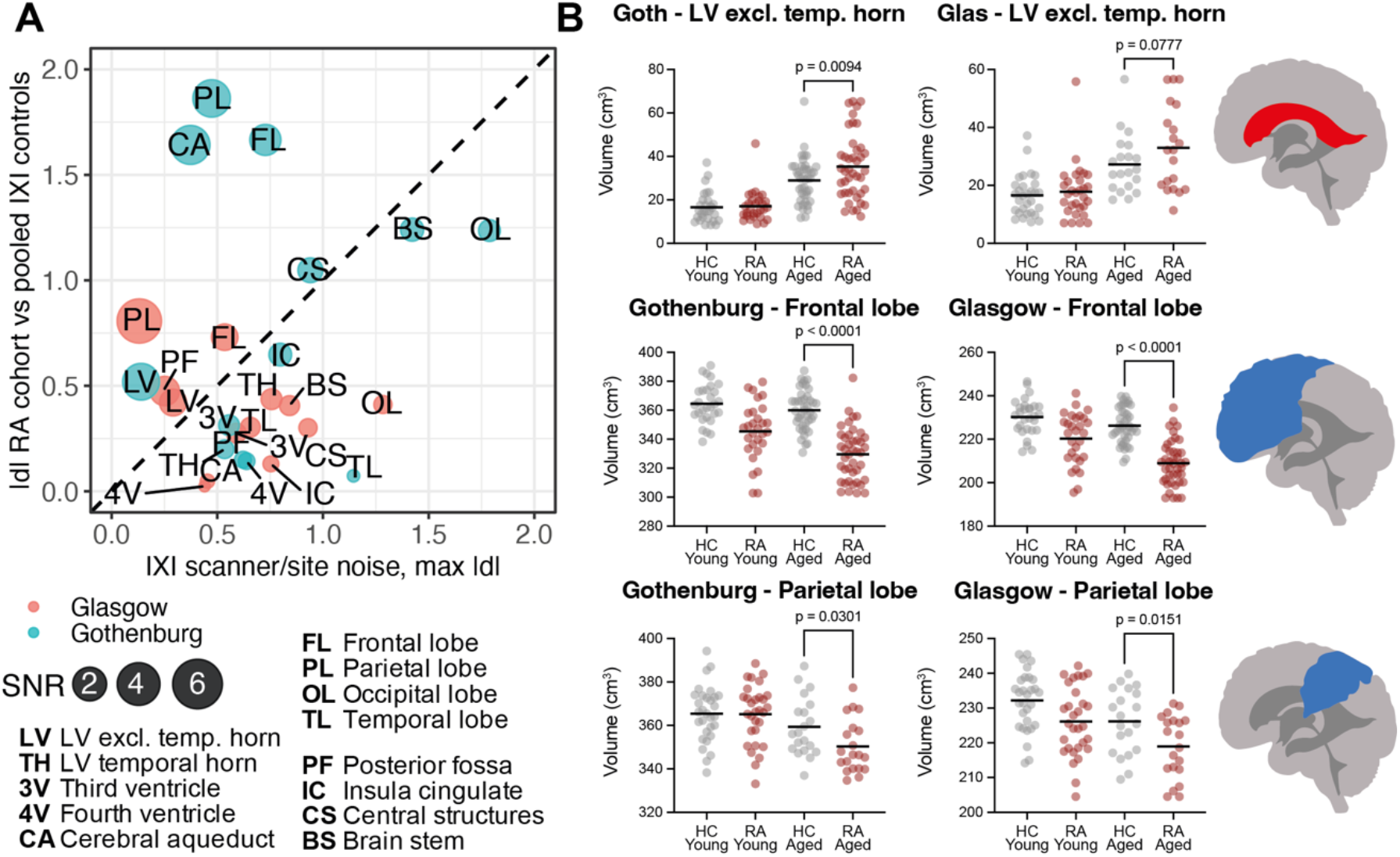
Rheumatoid arthritis is associated with reduced frontoparietal brain volumes and enlarged ventricles. Brain region volumes derived from magnetic resonance imaging (MRI) were analysed in two independent RA cohorts (Gothenburg, n = 71; Glasgow, n = 50) and age- and sex-matched healthy controls without diagnosed neurological disease from three IXI sites (HH, Guys, and IOP). Controls were strictly matched for age (maximum difference 3 years). **A)** Scatter plot of absolute effect sizes (|d|, Hedges’ g) for differences between RA patients and pooled IXI controls (y-axis) plotted against inter-site variability across IXI scanners (x-axis). Each point represents a brain region (abbreviations shown), colored by cohort and scaled by the signal-to-noise ratio (SNR = |d| / inter-site variability). The dashed diagonal line indicates equality between disease-related effects and scanner/site variability (|d| = noise). Points above the line represent regions where RA-associated differences exceed inter-site variability, whereas points below the line indicate effects smaller than scanner/site noise. **B)** Lateral ventricle (excluding temporal horn; upper panels), frontal lobe (middle panels) and parietal lobe (lower panels) volumes in the Gothenburg (left) and Glasgow (right) cohorts and corresponding healthy controls from the HH control cohort. Participants were stratified by age (<60 and ≥60 years). Values were winsorized at 2%. P-values were calculated using one-way ANOVA with post-hoc comparisons between older (≥60 years) RA and HC groups.

To further characterize the directionality and age dependence of the regional alterations identified in the signal-to-noise analysis, we performed age-stratified analyses focusing on brain regions with the most consistent disease-associated effects across cohorts (**Fig. 2B**). In both RA cohorts, patients older than 60 years demonstrated significantly enlarged lateral ventricles compared with age-matched healthy controls (Gothenburg: 34.5 versus 28.1 cm^3^, p = 0.004; Glasgow: 33.2 versus 25.6 cm^3^, p = 0.012). Also, older RA patients displayed reduced frontal lobe volumes, particularly in the Gothenburg cohort (330 versus 360 cm^3^, p < 0.0001), with similar patterns observed in the Glasgow cohort and in the parietal lobe across both cohorts. In contrast, corresponding regional differences were substantially weaker in patients younger than 60 years, consistent with the age-dependent increase in corrected brain-age gap observed in the brain-age analyses. Together, these findings indicate that the elevated brain-age gap in older RA patients is associated with a structural pattern characterized by ventricular enlargement and frontoparietal volume loss.

### RA patients display elevated serum biomarkers of neurodegeneration

To investigate whether the structural brain alterations observed in RA were accompanied by circulating biomarkers of neurodegeneration, we measured serum levels of brain-derived tau (BD-tau), glial fibrillary acidic protein (GFAP), and neurofilament light chain (NFL) in the Gothenburg RA cohort and age-matched healthy controls (**Fig. 3A**).

**Figure 3.**
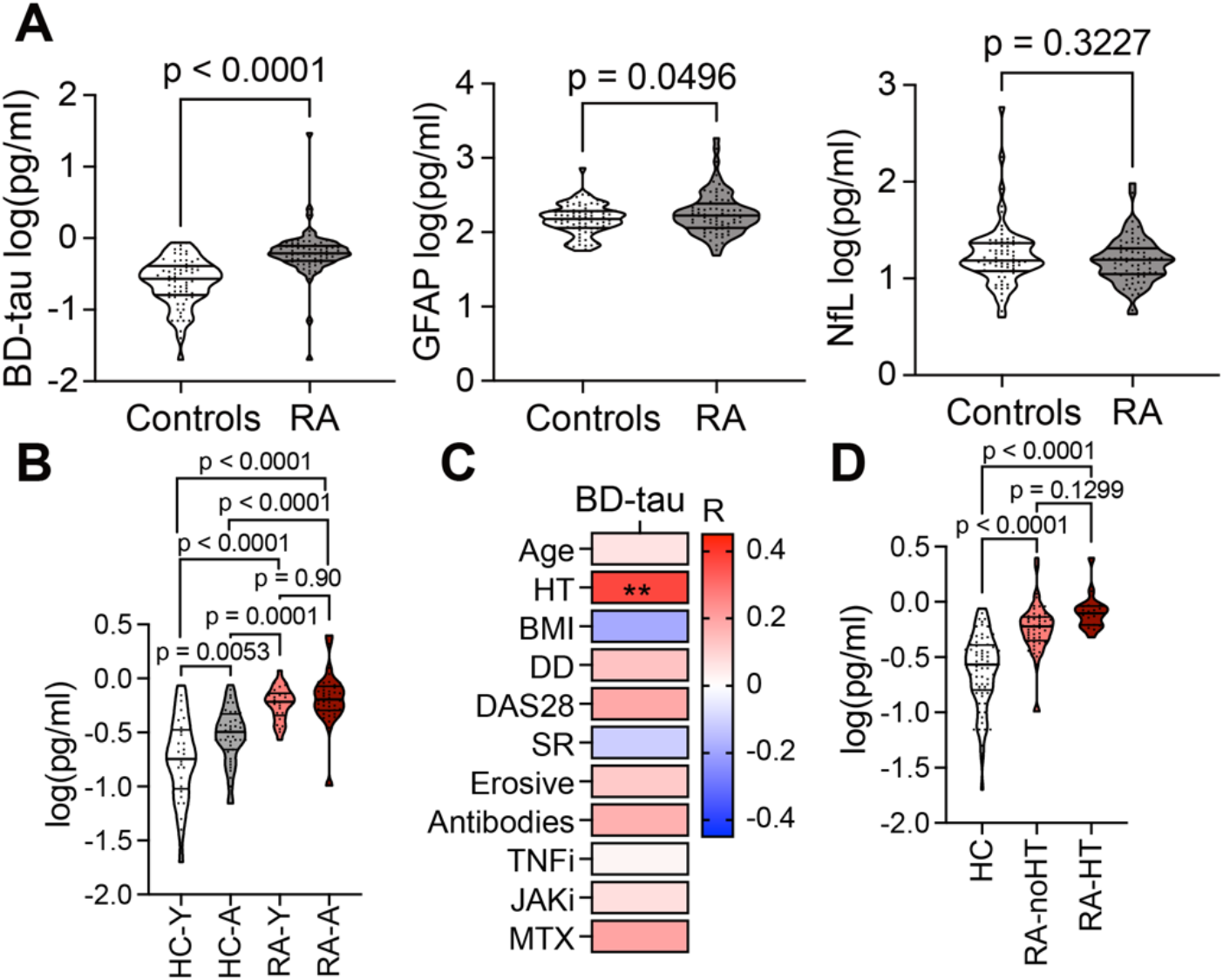
Patients with rheumatoid arthritis (RA) display elevated serum biomarkers of neurodegeneration. Serum levels of neurofilament light chain (NfL), brain-derived tau (BD-tau), and glial fibrillary acidic protein (GFAP) were measured in RA patients (Gothenburg cohort, n = 71) and age-matched healthy controls (HC). Control samples were pooled from three individuals per sample (71 pooled samples representing 213 individuals). **A)** Serum levels of BD-tau, GFAP, and NfL in RA versus HC. P-values were calculated using t-tests on logarithmically transformed data. **B)** Log-transformed and winsorized (3%) BD-tau levels in HC and RA patients stratified by age (<60 vs ≥60 years, left). P values were calculated using one-way ANOVA with post-hoc comparisons. **C)** Spearman correlation analysis between BD-tau (log-transformed and winsorized) and clinical variables (HT – hypertension, BMI – body-mass index, DD – disease duration, DAS28 – disease activity, SR – erythrocyte sedimentation rate, TNFi – tumour necrosis factor inhibitors, JAKi – Janus kinase inhibitors, MTX – methotrexate). The heatmap shows spearman R. **D)** Log-transformed and winsorized BD-tau levels in HC and RA patients stratified by hypertension status. P values were calculated using one-way ANOVA with post-hoc comparisons. *p < 0.05, **p < 0.01, ***p < 0.001, ****p < 0.0001.

RA patients displayed significantly elevated serum BD-tau levels compared with healthy controls (mean log-transformed BD-tau: 0.65 versus 0.31 pg/ml, p < 0.0001). GFAP levels were also modestly increased in RA (p = 0.0496), whereas NFL levels had no significant difference between groups (p = 0.32).

Because the MRI analyses demonstrated stronger age-associated neurostructural alterations in older RA patients, we next stratified BD-tau levels by age (<60 versus ≥60 years; **Fig. 3B**). BD-tau levels increased with age in both healthy controls and RA patients; however, RA patients exhibited substantially elevated BD-tau levels relative to controls in both age groups. The highest levels were observed in older RA patients (mean log-transformed BD-tau: 0.77 pg/ml versus 0.35 pg/ml in older healthy controls, p < 0.0001). Younger RA patients also showed elevated BD-tau levels compared with younger controls (0.62 versus 0.25 pg/ml, p = 0.0053). In contrast, BD-tau levels did not differ significantly between younger and older RA patients (p = 0.90).

Exploratory correlation analyses demonstrated that BD-tau levels were positively associated with hypertension status within the RA cohort (Spearman R = 0.31, p < 0.01; **Fig. 3C**), whereas correlations with disease duration, disease activity, inflammatory markers, and RA treatment were weak or absent. Stratification by hypertension status confirmed that RA patients with hypertension displayed the highest BD-tau levels (**Fig. 3D**). However, BD-tau levels also remained significantly elevated in RA patients without hypertension compared with healthy controls (p < 0.0001), indicating that the increase in BD-tau was not solely explained by hypertension.

### Association of myeloid transcriptional signatures with brain-age gap in RA

To investigate whether brain aging in RA was associated with specific myeloid transcriptional programs, we quantified published transcriptional signatures representing aged monocytes, disease-associated microglia (DAM), and disease-inflammatory macrophages (DIM) in peripheral CD14+ monocytes from the Gothenburg RA cohort using ssGSEA (**Fig. 4**). The aged monocyte signature was derived from the study by Shchukina et al. ^22^, which compared circulating monocytes of healthy young and aged individuals. The DAM and DIM signatures were derived from human adult differentially expressed gene sets reported by Silvin et al. ^13^, generated from an integrated single-cell myeloid atlas (“M-Verse”) of aging and neurodegeneration-associated CNS macrophage populations.

**Figure 4.**
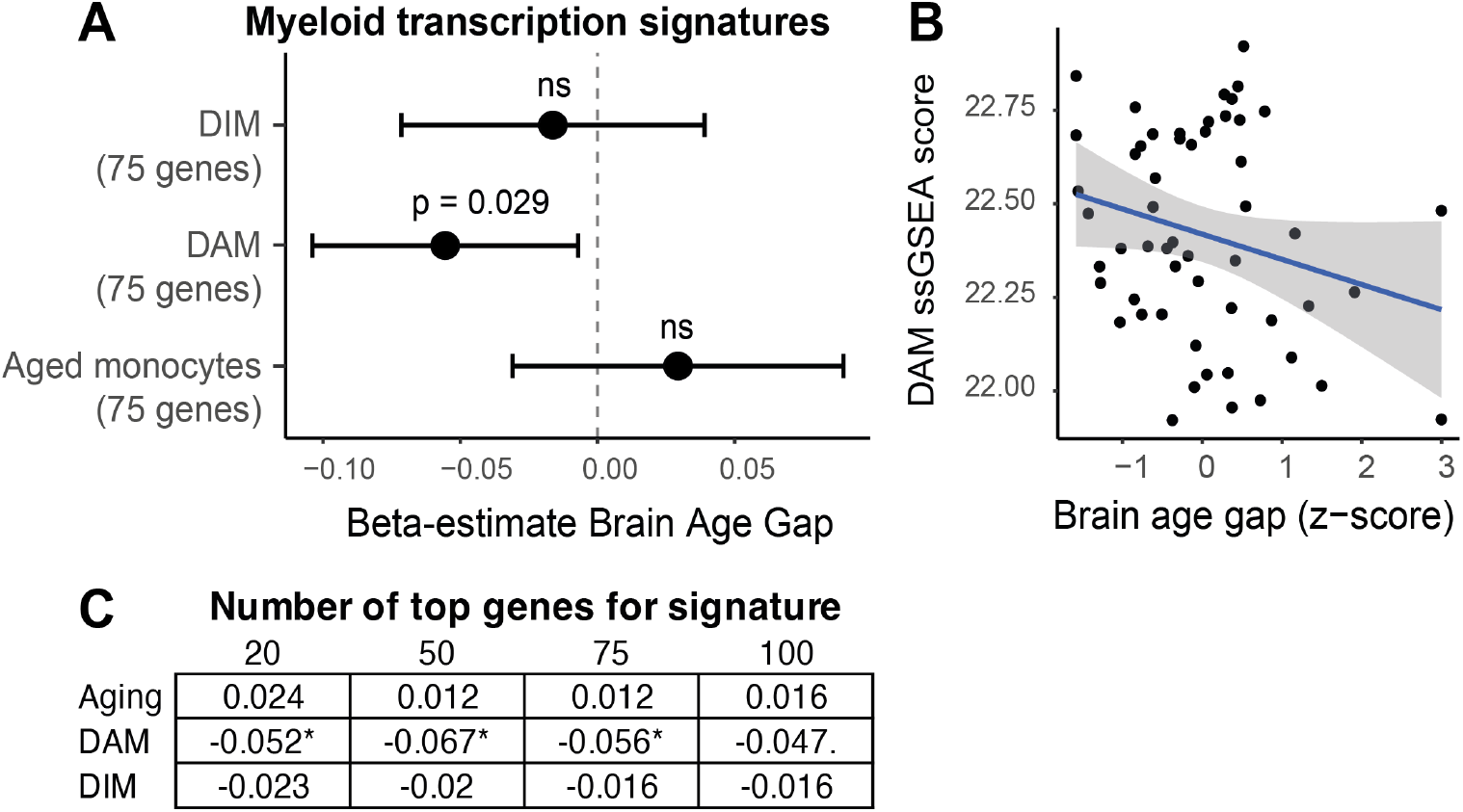
Association of myeloid transcriptional signatures with brain age gap in rheumatoid arthritis (RA). CD14+ monocytes were isolated from peripheral blood of patients with RA (Gothenburg cohort, n = 54), stimulated with LPS for 2 h, and analysed using RNA sequencing. Published myeloid transcriptional signatures representing aged monocytes ^22^, disease-associated microglia (DAM)^13^, and disease-inflammatory macrophages (DIM) ^13^ were quantified using single-sample gene set enrichment analysis (ssGSEA). Associations between signature scores and the difference between MRI-predicted brain age and chronological age (brain age gap) were assessed after winsorisation of the brain age gap variable at 2%. **A)** Adjusted beta estimates (±95% confidence intervals) for the association between brain age gap and monocyte signatures. Each signature consisted of the top 75 positively associated genes. Associations were assessed using linear regression models adjusted for chronological age and sequencing batch. **B)** Scatterplot showing the association between the DAM ssGSEA score and brain age gap (z-score). The solid line represents the linear regression fit and the shaded area indicates the 95% confidence interval. **C)** Robustness analysis across different signature sizes (top 20, 50, 75, and 100 genes). Values indicate adjusted beta estimates from linear regression models adjusted for chronological age and sequencing batch.

No significant association was observed between brain-age gap and the aged monocyte signature (β = 0.029, p = 0.345) or the DIM signature (β = −0.016, p = 0.566) after adjustment for chronological age and sequencing batch (Figure 4A). In contrast, the DAM signature showed a significant inverse association with brain-age gap (β = −0.056, p = 0.029), indicating lower enrichment of the DAM-related transcriptional program in patients with greater MRI-predicted brain aging. The association between DAM score and brain-age gap is shown in **Fig. 4B**.

To assess robustness of the findings, signature analysis was repeated using different signature sizes (top 20, 50, 75, and 100 positively associated genes from each reference dataset). The inverse association between the DAM signature and brain age gap remained directionally consistent across all tested gene-set sizes, whereas aged monocyte and DIM signatures showed no stable associations (**Fig. 4C**).

## Discussion

Using two independent cohorts of RA patients, this study demonstrated that RA disease was associated with volume reduction in cortical brain regions of the frontoparietal lobes, higher levels of BD-tau in serum and the enlarged lateral ventricles in aged patients. Our findings elaborate on a previous finding of a higher ventricle-to-brain area ratio measured in a single cross-section of brain in RA patients with disease duration over 15 years ^23^. In neurodegenerative disease, accelerated ventricle enlargement is generally associated with cognitive impairment. Indeed, RA patients have repeatedly been reported to perform significantly worse than controls in cognitive tests ^24-28^. A recent meta-analysis revealed cognitive impairment in RA patients, who were scored 1.27 points lower than controls on the Montreal Cognitive Assessment ^27^. In line with this, genome-wide association studies suggested a link between RA and frontotemporal dementia, with accelerated cerebral atrophy centred on the frontal lobe cortex, and a shared single nucleotide polymorphism in the human leukocyte antigen gene ^29^. Additionally, RA was identified as an independent risk factor for developing dementia ^30^, suggesting shared molecular pathology behind these conditions. Despite the presence of several common features between RA and dementia patients, the structural changes in brain that we observed in aging RA patients were not identical to those in Alzheimer’s disease. The regions found to be small in RA patients overlapped with regions affected at later stages of Alzheimer’s disease ^31^, but the amygdala, hippocampus, entorhinal cortex and para-hippocampal cortex regions typically affected at earlier stages of Alzheimer’s disease were not reduced in RA patients on a cohort level.

The elevated circulating BD-tau levels observed in RA patients further support the presence of neuroaxonal alterations associated with RA. BD-tau has emerged as a relatively CNS-enriched marker associated with cortical neurodegeneration and cognitive decline in Alzheimer’s disease and related disorders ^7^. In the present study, BD-tau elevation was observed in both younger and older RA patients and was not clearly associated with RA disease duration, suggesting that neuroaxonal alterations may occur relatively early during RA or reflect mechanisms not solely dependent on cumulative inflammatory burden. In contrast, serum NFL levels were not significantly elevated in RA, potentially indicating that the observed neurostructural alterations were more strongly related to cortical neuronal injury than to widespread subcortical axonal degeneration. This interpretation is consistent with the predominant reduction of frontoparietal cortical volume observed in the MRI analyses.

In the present study, peripheral CD14+ monocytes from RA patients with increased brain-age gap showed lower enrichment of a DAM-related transcriptional signature derived from human CNS macrophage populations associated with aging and neurodegeneration. Importantly, blood monocytes are generally not equivalent to brain-resident microglia, and the observed transcriptional overlap should therefore be interpreted cautiously. However, monocyte-derived macrophages can infiltrate the CNS under inflammatory conditions, and peripheral myeloid cells may partially reflect systemic immune programs relevant to neuroinflammatory processes. Notably, DAM phenotype in neurodegenerative disease is thought to have context-dependent function, including both pro-inflammatory activity and a potentially protective role related to debris clearance and tissue remodelling. The inverse association between DAM signature enrichment and brain-age gap observed in this study may therefore suggest impaired activation of protective myeloid programs in patients with more advanced neurostructural alterations. In parallel, RA patients demonstrated elevated circulating GFAP levels, consistent with increased astrocytic activation or injury. Together, these findings support the possibility that systemic immune dysregulation in RA is possibly accompanied by neuroimmune alterations involving both myeloid and astrocytic components. Furthermore, previous studies demonstrated an enrichment of IL-1b in cerebrospinal fluid compared to serum in RA patients ^32^, and an association between high cortical microglia density and genetic predisposition for RA ^33^.

An alternative biological mechanism that would facilitate brain atrophy in RA is cardiovascular morbidity, which can promote atrophy by restricting the blood supply of brain tissue. Cardiovascular disease is highly prevalent in RA patients ^34^. We found that hypertension status was associated with higher brain age gap and serum levels of BD-tau. However, hypertension status could not explain all differences in brain tissue volumes between RA patients and controls.

This study has several limitations. First, the cross-sectional design does not allow conclusions regarding longitudinal progression of brain aging in RA. Second, healthy controls were drawn from an external multi-site imaging dataset rather than prospectively recruited alongside the RA cohorts, and residual scanner- or site-related effects therefore cannot be fully excluded despite replication across independent cohorts and explicit modelling of inter-site variability. An important next step will be longitudinal investigation of brain aging trajectories in RA across different age groups. Finally, the treatment analyses were exploratory and should be interpreted cautiously because treatment allocation and indication are inherently confounded in observational studies. To determine whether TNF inhibition in combination with methotrexate attenuates age-related neurostructural decline in RA, randomized controlled studies are required.

In summary, our findings provide converging evidence that RA is associated with structural brain alterations consistent with advanced brain aging, particularly in older individuals. Across independent cohorts, RA patients demonstrated increased brain-age gap, ventricular enlargement, reduced frontoparietal brain volumes, and elevated circulating markers of neurodegeneration. The age dependence of these findings, together with the association between structural alterations and immunomodulatory treatment, supports a link between chronic systemic inflammation and the accelerated age-related neurostructural decline in RA. Prospective controlled studies will determine whether RA is associated with accelerated rates of brain tissue loss over time, identify critical periods of vulnerability during aging, and clarify whether immunomodulatory therapies modify the progression of neurostructural decline. Integration of longitudinal MRI, blood biomarkers, cognitive assessments, and immune profiling may advance understanding the mechanisms linking systemic inflammation to brain aging in RA.

## Materials and Methods

### Study participants

This study is based on two independent cohorts of RA patients: the Gothenburg and Glasgow cohorts. Patient characteristics are given in **Table 1**.

### Gothenburg cohort

Patients with established RA (female, n=71), meeting the 2010 American College of Rheumatology/European League Against Rheumatism classification criteria^35^, were recruited from the Rheumatology Clinic at Sahlgrenska University Hospital from 2019 to 2020. The patients were examined by an experienced rheumatologist, underwent venous blood sampling, completed a battery of clinical questionnaires, and had a magnetic resonance (MR) imaging scan of the cranium. All patients gave written informed consent prior to participating in the study. The study was approved by the Swedish Ethical Review Authority (Dnr. 2019-03787). The study is registered at ClinicalTrials.gov (ID NCT03449589). At the time of sample collection, 14 patients were treated with methotrexate monotherapy, ten patients combined methotrexate with TNF inhibitors (TNFi), twelve patients with JAK inhibitors (JAKi), nine with other synthetic or biologic anti-rheumatic drugs, and seven patients had no anti-rheumatic treatment. In total, five patients used oral corticosteroids.

### Glasgow cohort

Patients with established RA (76% female, n=50) were recruited from regional rheumatology services in the United Kingdom. All participants met the 2010 American College of Rheumatology/European League Against Rheumatism classification criteria ^35^. Patients were excluded if they had MRI contraindications such as metal implants or were left-handed. The patients underwent multimodal brain imaging, venous blood sampling, and a battery of clinical questionnaires. All participants gave written informed consent following the Declaration of Helsinki. The study was approved by the North of Scotland Research Ethics Committee.

written informed consent following the Declaration of Helsinki. The study was approved by the North of Scotland Research Ethics Committee.

### Controls

To compare changes in the brain tissue of RA patients with healthy individuals, we used the IXI cohort (http://brain-development.org/ixi-dataset/), which includes MR images of 245 women (mean age in years=46.8, SD=16.5, range=20.1-86.2) and 336 men (mean age in years=50.11, SD=16.4, range=20.0-86.3) with no history of RA or neurological disease.

### MRI and image analysis

#### Gothenburg cohort

MR images were acquired using a 3-tesla scanner with a 32-channel SENSE head coil (Philips Gyroscan Achieva 3T; Philips Healthcare). T1-weighted images were acquired with the following parameters: flip angle = 8°, echo time = 4.0 ms, repetition time = 8.4 ms, SENSE factor = 2.7, turbo field echo factor = 240 and voxel size = 1 × 1 × 1 mm^3^.

#### Glasgow cohort

The MRI sequence included a T1-weighted fast-field echo 3D structural sequence. T1-weighted images were acquired with the following parameters: flip angle = 8°, echo time = 3.8 ms, repetition time = 8.2 ms, inversion time = 1018 ms, voxel size = 0.94 × 0.94 × 1 mm^3^, matrix size 240 × 240 with 160 slices and field of vision = 240 mm.

Images were pre-processed with field inhomogeneity correction (N4ITK) software. Morphometric analysis was carried out on T1-weighted MRI images using ensemble machine learning models for semantic image segmentation. Brain masking (skull stripping) was performed with Pincram ^36^. MAPER ^21^ was used to label and measure brain regions. For training data, we used the 30 cranial MR images of healthy volunteers contained in the Hammers Adult Brain Atlas Database ^37-40^ along with reference label sets manually produced by experts into 120 regions (cortical as well as subcortical; pre-release version). MAPER processing includes three-class probability mapping (seg_EM from NiftySeg, https://github.com/KCL-BMEIS/NiftySeg, considering grey matter (GM), white matter (WM), and cerebrospinal fluid); crisp label images that distinguish these tissue classes were generated from the resulting maps. For each left-right paired region, we calculated the sum of the right and left volumes. In total, the volumes of 62 brain regions were analysed (**Supplementary table 1**). We also ran analyses in which brain regions were combined into super regions (brainstem, central structures, frontal lobe, insula and cingulate, occipital lobe, posterior fossa, temporal lobe, and ventricles (**Supplementary table 1**)). All volumes were normalized by intracranial volume as determined by Pincram and scaled by the mean intracranial volume of the IXI cohort. Please note that the images captured by this methodology provide structural information only.

### Cell isolation and culture

The samples were collected between 7 and 10 a.m. after overnight fasting. For serum preparation, the blood sample was obtained from the cubital vein into vacuum containers (BD Vacutainer). Serum samples and culture supernatants were stored at -70°C until use. Human peripheral blood mononuclear cells were separated using density gradient centrifugation with Lymphoprep (Axis-Shield PoC As, Norway). CD14+ cells were isolated via positive selection (Stemcell Technologies 17858) from the mononuclear cell mixture. Flow cytometry analysis of isolated cells demonstrated a CD14+ cell purity of 78-90%. The cells were plated (1.25 × 10^6^ cells/ml) and stimulated for 2 hours with 5 μg/ml lipopolysaccharide (LPS, Sigma-Aldrich, St. Louis, Missouri, USA) in RPMI medium supplemented with 5% foetal bovine serum (Sigma-Aldrich), 2 mM Glutamax (Gibco, Waltham, Massachusetts, USA), 50 μg/ml gentamicin (Sanofi-Aventis, Paris, France) and 50 μM β-mercaptoethanol (Gibco) at 37°C in a humidified 5% CO_2_ incubator. Culture supernatants were collected for the analysis of cell products by ELISA.

### RNA sequencing and analysis

RNA was extracted from CD14+ cell cultures using the Total RNA Purification Micro Kit (Norgen Biotek Corp, Thorold, Ontario, Canada). RNA quality was assessed with an RNA 6000 Pico Kit on the 2100 Bioanalyzer System (Agilent Technologies, Inc., Santa Clara, CA, USA). Sequencing was performed using RNA-Seq on a Hiseq 2000 System (Illumina, Inc., San Diego, CA, USA) at the BEA Core Facility, Karolinska Institutet, Sweden.

Transcriptomic analyses were performed in R (R Foundation for Statistical Computing, Vienna, Austria) using the limma (v3.56.2) and GSVA (v1.48.3) packages. Analyses were restricted to protein-coding genes. Genes were retained for analysis if expression exceeded log2(6) in at least 20% of samples.

Associations between monocyte gene expression and brain age gap were analysed using robust linear models including brain age gap, chronological age, and sequencing batch as covariates. Brain age gap values were winsorized at 2% prior to analysis to reduce the influence of extreme values. Empirical Bayes moderation was applied using the eBayes function in limma, and p-values were adjusted for multiple testing using the Benjamini–Hochberg method.

Published myeloid transcriptional signatures representing aged monocytes, disease-associated microglia (DAM), and disease-inflammatory macrophages (DIM) ^13,22^ were quantified using single-sample gene set enrichment analysis (ssGSEA) implemented in the GSVA package. Signature scores were calculated using the top positively associated genes from each reference dataset and tested for association with brain age gap using linear regression models adjusted for chronological age and sequencing batch.

### Auto-antibody measurement

Rheumatoid factor (RF) and antibodies against cyclic citrullinated proteins (anti-CCP) were measured in serum samples by the accredited Clinical Immunology Laboratory at Sahlgrenska University Hospital. Anti-CCP was measured using an automated multiplex method (BioPlex 2200 System, Bio-Rad Laboratories AB, Solna, Sweden). Levels of anti-CCP above 3.0 U/ml were considered positive. Antibodies against RF were measured by Dual Detection Rate Nephelometric Technology (Immage 800, Beckman Coulter AB, Solna, Sweden). Levels of RF over 20 U/ml were considered positive.

### Measurements of NFL, GFAP and brain-derived tau

Serum levels of NFL and GFAP were measured using a commercially available duplex Single molecule array (Simoa) assay, according to instructions by the kit manufacturer (Quanterix, Billerica, MA, USA). Brain-derived tau concentration was measured using an in-house Simoa assay, as previously described^7,8^. For the analysis, a control group was created by pooling equal volumes of 3 sex and age-matched RA-free controls per each RA patient into one control sample. Calibrators were run in duplicates and obvious outlier calibrator replicates were masked before curve fitting. Two quality control (QC) samples were run in duplicates in the beginning and the end of each run. **NFL:** For a sample with a concentration of 13.4 pg/mL, repeatability was 9.4% and intermediate precision was 9.4%. For a QC sample with a concentration of 105 pg/mL, repeatability was 11.2% and intermediate precision was 18.1%. Validated Measurement Range = 2.7 - 1580 pg/mL. **GFAP:** For a QC sample with a concentration of 67.0 pg/mL, repeatability was 8.2% and intermediate precision was 17.8%. Validated Measurement Range = 40 - 29400 pg/mL. **Brain-derived tau:** For a QC sample with a concentration of 1.5 pg/mL, repeatability was 1.2% and intermediate precision was 18.2%. For a QC sample with a concentration of 5.7 pg/mL, repeatability was 5.3% and intermediate precision was 11.5%. Validated Measurement Range: 0.062 - 16 pg/mL.

### Cardiovascular risk assessment

Cardiovascular risk was estimated using the Framingham risk score (FRS) ^41^. To calculate the FRS, we used the following cardiovascular risk factors: high density lipids (HDL), systolic blood pressure (SBP), age, smoking, diabetes, and total cholesterol. The 10-year risk score was estimated using the online calculator provided by the Framingham Heart Study, available at https://www.framinghamheartstudy.org/fhs-risk-functions/cardiovascular-disease-10-year-risk/. Spearman correlation analysis was used to calculate the association between FRS and brain region volumes. Multiple linear regression analysis was used to investigate the association between brain region volumes and individual cardiovascular risk factors. FRS cardiovascular risk factors were standardized in Prism for MacOS (GraphPad Software, Boston, MA, USA; version 10.3.1) software to facilitate visualization.

### Statistical analysis

#### Brain-age prediction analysis

A brain-age prediction model was trained using magnetic resonance imaging (MRI)-derived regional brain volumes from healthy controls in the IXI dataset (n = 566; HH, Guys, and IOP sites pooled). Regional brain volumes were normalized to intracranial volume (ICV) and converted from mm^3^ to cm^3^ prior to analysis. Brain stem volume was excluded because this region was sensitive to differences in MRI field-of-view across scanners. The following regional volumes were included as predictors in the brain-age model: frontal lobe, parietal lobe, temporal lobe, occipital lobe, insula/cingulate region, central structures, posterior fossa, lateral ventricles excluding temporal horns, temporal horns of the lateral ventricles, third ventricle, and fourth ventricle.

Chronological age was predicted using multiple linear regression with regional brain volumes as independent variables. To facilitate interpretation of regional contributions to age prediction, a second model using z-score standardized variables was generated and standardized β-estimates were extracted.

The trained model was subsequently applied to healthy IXI controls and RA patients from the Gothenburg and Glasgow cohorts to estimate MRI-predicted brain age. As commonly observed in brain-age prediction models, raw predicted age showed age-dependent bias, with younger individuals tending to have their age overestimated and older individuals tending to have their age underestimated. To account for this bias, the relationship between predicted age and chronological age was modelled in the IXI healthy controls using natural splines with three degrees of freedom. This model was used to estimate the expected predicted age for a healthy individual at each chronological age.

Corrected brain-age gap was defined as the difference between an individual’s predicted brain age and the expected predicted age for a healthy control of the same chronological age. Following correction, brain-age gap was no longer associated with chronological age in the IXI control cohort. Positive corrected brain-age gap values indicated an older-appearing brain than expected for age, whereas negative values indicated a younger-appearing brain.**Statistical analysis of brain-age gap**. The primary hypothesis tested whether RA patients displayed a positive corrected brain-age gap relative to the healthy reference population. This was assessed using a one-sample t-test against zero. Associations between corrected brain-age gap and chronological age in RA patients were assessed using linear regression models. To determine whether age-related changes in corrected brain-age gap differed between RA patients and healthy controls, linear regression models including a Group × Age interaction term were fitted using pooled IXI controls and RA patients. Cohort consistency between the Gothenburg and Glasgow RA cohorts was evaluated using Cohort × Age interaction models. For visualization purposes, RA patients were additionally stratified into younger (<60 years) and older (≥60 years) age groups. Non-parametric Wilcoxon tests were performed as sensitivity analyses for age-stratified comparisons.

### Supplementary validation analyses

Brain-age model performance in healthy IXI controls was assessed using Pearson correlation coefficients and coefficients of determination (R^2^) between chronological age and predicted brain age. Mean absolute error (MAE) was calculated as the absolute difference between predicted and chronological age averaged across all individuals. To confirm successful bias correction, associations between corrected brain-age gap and chronological age in IXI controls were assessed using Pearson correlation analysis and linear regression. To evaluate robustness across MRI scanner sites, leave-one-site-out validation was performed within the IXI dataset. For each iteration, the brain-age model and bias-correction model were trained using two IXI sites and applied to the remaining left-out site. Corrected brain-age gap values in the left-out site were subsequently assessed for deviation from zero.

### Scanner/site robustness analysis of regional brain volume differences

To evaluate whether disease-associated regional brain volume differences exceeded variability attributable to MRI scanner/site effects, a robustness analysis was performed using healthy controls from the IXI dataset and the independent RA cohorts from Glasgow and Gothenburg. Analyses were performed separately for each RA cohort. For each RA cohort, healthy IXI controls were matched 1:1 to RA participants by sex and chronological age within a strict maximum age difference of 3 years. Controls from the three IXI MRI sites (Guys, HH, and IOP) were pooled for matching. Because the IOP site contained fewer age-matched individuals in older age groups, some IXI controls were reused across matched comparisons. MRI-derived regional brain volumes were grouped into combined super-regions, including frontal, parietal, temporal, and occipital lobes, insula/cingulate region, central structures, posterior fossa, and cerebrospinal fluid compartments. Scanner/site variability within the IXI healthy control dataset was estimated by calculating pairwise standardized effect sizes between IXI MRI sites (Guys, HH, and IOP) for each regional volume. For each region, the maximum absolute standardized mean difference across all pairwise site comparisons were retained as the estimate of scanner/site noise. Standardized effect sizes were calculated using Hedges’ g. Disease-associated effects were estimated by comparing each RA cohort with pooled matched IXI healthy controls using Hedges’ g for each regional volume. Absolute effect sizes (|d|) were used to quantify the magnitude of disease-associated differences independently of directionality. Confidence intervals for effect sizes were estimated analytically from the pooled standard error of the standardized mean difference. A signal-to-noise ratio (SNR) was subsequently calculated for each region as:|*d*| *(RA versus IXI) / scanner-site noise*. To avoid inflation of SNR values in regions with minimal inter-site variability, a minimum scanner/site noise floor of 0.1 was applied. Scatter plots were generated with disease-associated effect sizes on the y-axis and scanner/site variability on the x-axis.

### Outliers

Extreme values were winsorized prior to association analyses with clinical variables to reduce the influence of highly influential observations. Corrected brain-age gap values were winsorized at the 2nd and 98th percentiles, and BD-tau values at the 3rd and 97th percentiles.

### Use of Artificial Intelligence Generated Content tools Spelling and grammar were improved with ChatGPT. Data and materials availability

RNA sequencing data from human CD14+ cells are deposited in Gene Expression Omnibus at the National Centre of Biotechnology Information (NCBI) with the accession code GSE201670 ^42^. Due to ethical and legal restrictions related to patient confidentiality and the handling of pseudonymized MRI data under the EU General Data Protection Regulation, the imaging and associated clinical datasets are not publicly available. Data may be made available upon reasonable request to the corresponding author, subject to review of a research proposal, relevant ethical approval, and a data use agreement to ensure compliance with data protection and ethical requirements.

## Acknowledgments

We are thankful to the research nurses Anneli Lund and Marie-Louise Andersson, Rheumatology Clinic, Sahlgrenska University Hospital, for assistance with blood sampling. We thank all the RA patients who participated in this study.

This work has been funded by grants from the Swedish Research Council (CW: 2020-00592 and 2025-02120, MB: 2017-03025 and 2017-00359), the FOREUM Foundation for Research in Rheumatology (0057/2025), the Swedish Rheumatism Association (CW: R-982397, R-995855 and R-1013796; MB: R-566961, R-751351 and R-860371; RP: R-969562, R-862061, R-995509), the King Gustaf V:s 80-year Foundation (CW: FAI-2024-1119, MB: FAI-2018-0519, FAI-2020-0653 and FAI-2022-0882), the Regional agreement on medical training and clinical research in the Western Götaland county (MB: ALFGBG-717681 and ALFGBG-965623; RP: ALFGBG-965012 and ALFGBG-926621), the University of Gothenburg (CW: GU2022/1937), Inger Bendix Foundation (CW: #87) and Tore Nilson Foundation (CW: 2019-00742 and 2024-216) and Pfizer IIR award (NB). MR image processing was enabled by resources provided by the National Academic Infrastructure for Supercomputing in Sweden (NAISS), partially funded by the Swedish Research Council through grant agreement no. 2022-06725. HZ is a Wallenberg Scholar and a Distinguished Professor at the Swedish Research Council supported by grants from the Swedish Research Council (#2023-00356, #2022-01018 and #2019-02397), the European Union’s Horizon Europe research and innovation programme under grant agreement No 101053962, Swedish State Support for Clinical Research (#ALFGBG-71320), the Alzheimer Drug Discovery Foundation (ADDF), USA (#201809-2016862), the AD Strategic Fund and the Alzheimer’s Association (#ADSF-21-831376-C, #ADSF-21-831381-C, #ADSF-21-831377-C, and #ADSF-24-1284328-C), the European Partnership on Metrology, co-financed from the European Union’s Horizon Europe Research and Innovation Programme and by the Participating States (NEuroBioStand, #22HLT07), the Bluefield Project, Cure Alzheimer’s Fund, the Olav Thon Foundation, the Erling-Persson Family Foundation, Familjen Rönströms Stiftelse, Stiftelsen för Gamla Tjänarinnor, Hjärnfonden, Sweden (#FO2022-0270), the European Union’s Horizon 2020 research and innovation programme under the Marie Skłodowska-Curie grant agreement No 860197 (MIRIADE), the European Union Joint Programme – Neurodegenerative Disease Research (JPND2021-00694), the National Institute for Health and Care Research University College London Hospitals Biomedical Research Centre, the UK Dementia Research Institute at UCL (UKDRI-1003), and an anonymous donor.

## Author Contributions

Conceptualization: MIB, RH

Methodology: MCE, KE, KS, AG, MH, RP, HZ, KB, NB, STS

Visualization: CW Supervision: MIB and RH

Writing—original draft: CW

Writing—review & editing: MCE, KE, KS, AG, MH, RP, HZ, KB, NB, STS, MIB, RH.

## Supplementary material

**Figure S1.**
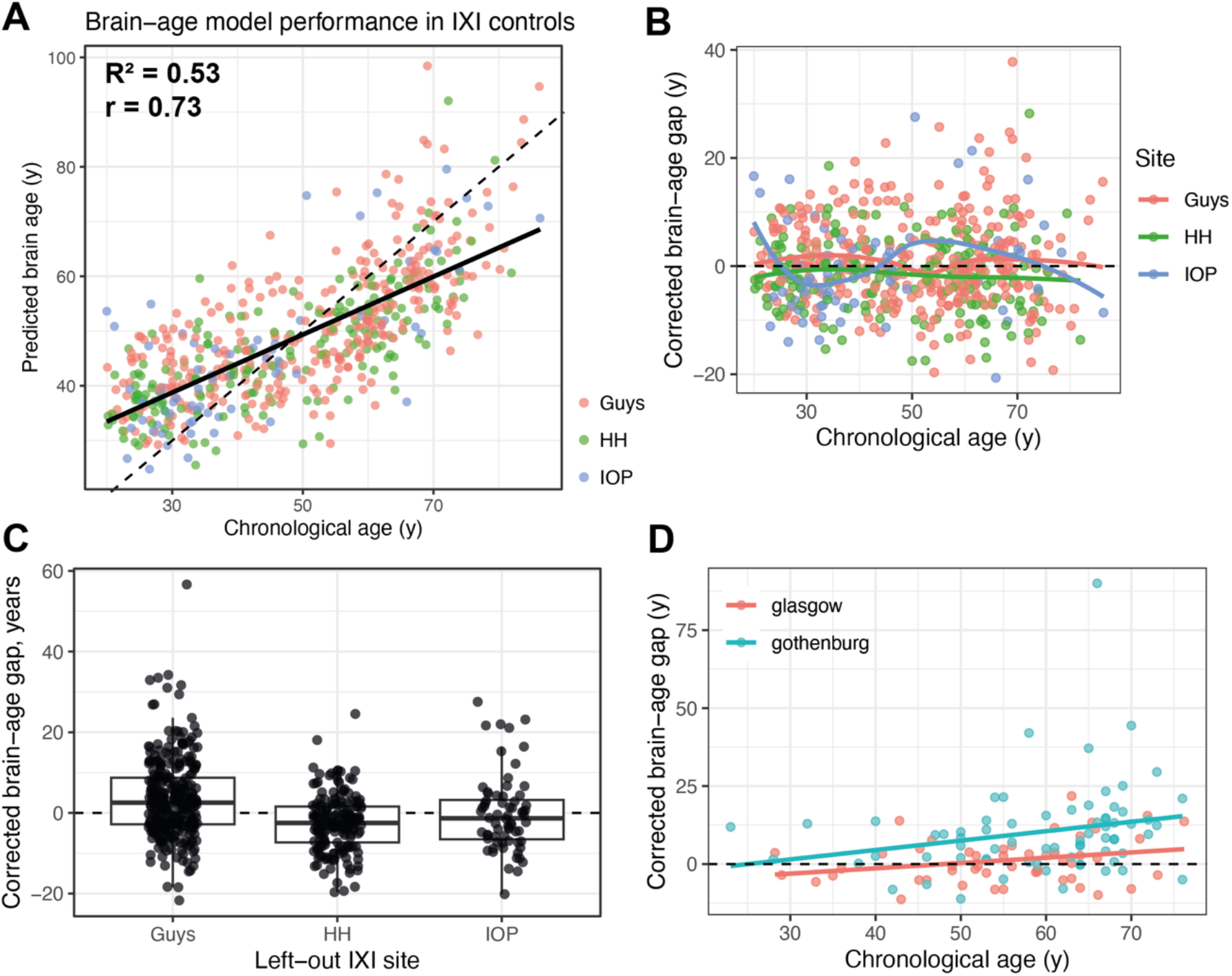
Validation of the MRI-derived brain-age prediction model in healthy IXI controls. **A)** Predicted brain age versus chronological age in healthy controls from the IXI dataset (HH, Guys, and IOP sites pooled; n = 566). Brain age was predicted using a linear regression model trained on intracranial volume-normalized regional brain volumes. The solid black line indicates the fitted linear regression model and the dashed line represents the line of identity (predicted age = chronological age). Model performance metrics are shown in the panel (R^2^, Pearson correlation coefficient r). **B)** Corrected brain-age gap in IXI healthy controls plotted against chronological age following bias correction using natural splines. Corrected brain-age gap was calculated as predicted brain age minus the expected predicted age for a given chronological age estimated from the IXI reference population. The absence of a systematic age-dependent trend after correction indicates successful removal of age-related prediction bias. Coloured lines represent site-specific smoothing curves for the HH, Guys, and IOP sites. **C)** Leave-one-site-out validation of the brain-age model across IXI MRI sites. For each analysis, the brain-age model and bias correction model were trained using two IXI sites and applied to the remaining left-out site. Boxplots and individual data points show corrected brain-age gap values for each left-out site. Dashed horizontal lines indicate zero corrected brain-age gap. **D)** Corrected brain-age gap plotted against chronological age separately for the Gothenburg and Glasgow RA cohorts.

**Figure S2.**
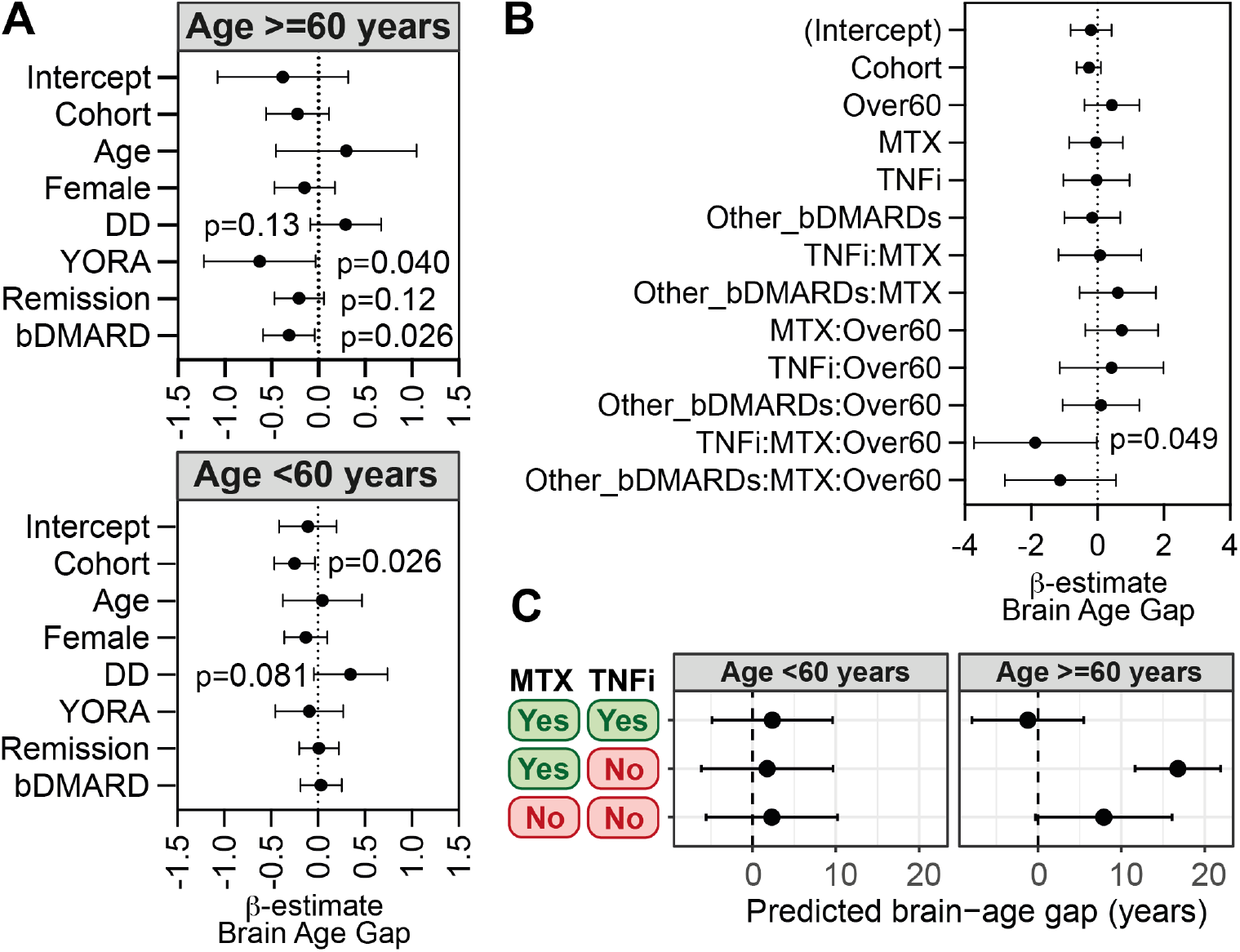
Association between rheumatoid arthritis (RA) clinical and treatment variables and brain-age gap. Pooled analysis of clinical and treatment variables in the Gothenburg (n=71) and Glasgow (n=50) cohorts of patients with established RA. **A)** Linear regression analyses stratified by age group (<60 years and ≥60 years) showing associations between clinical variables and brain-age gap. β-estimates and 95% confidence intervals are shown. Dashed vertical lines indicate no effect (β = 0). P-values are shown for variables reaching nominal significance or trend level (p<0.15). independent variables included cohort, age, sex, disease duration (DD), young-onset RA (YORA), remission status, and biologic disease-modifying anti-rheumatic drug (bDMARD) treatment. **B)** Multivariable linear regression model examining associations between methotrexate (MTX), TNF inhibitor (TNFi), other biologic DMARDs (Other_bDMARDs), age group (Over60), and their interaction terms with standardized brain-age gap. β-estimates and 95% confidence intervals are shown. **C)** Estimated marginal means from the regression model in panel B showing predicted brain-age gap across treatment groups stratified by age (<60 years and ≥60 years). Predicted values with 95% confidence intervals are shown for patients receiving no MTX/no TNFi, MTX without TNFi, and combined MTX+TNFi treatment, while holding Other_bDMARDs = 0 and Cohort = 0. Brain-age gap values are presented in years.

**Figure S3.**
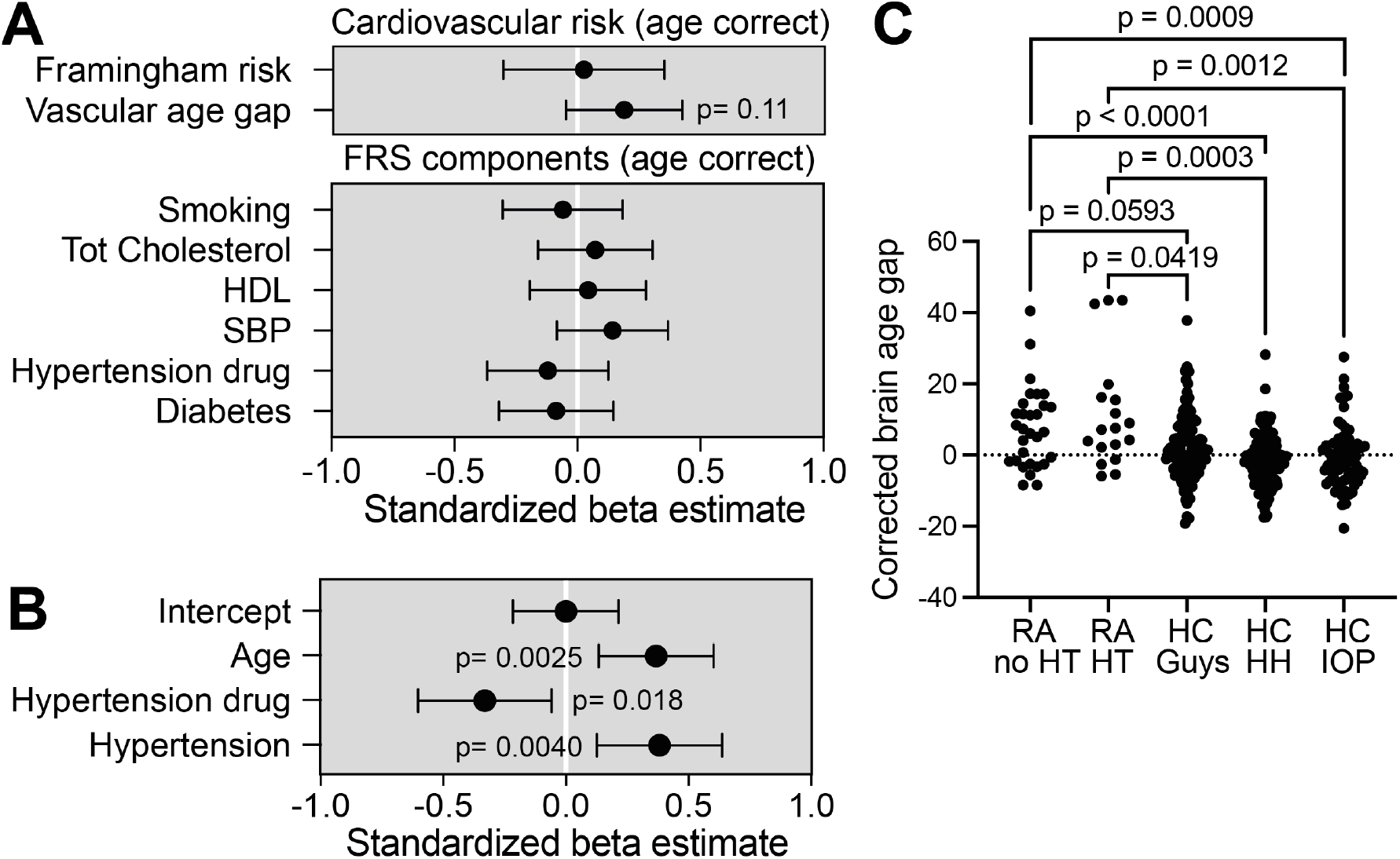
Cardiovascular risk and corrected brain-age gap in rheumatoid arthritis (RA). **A)** Age-adjusted linear regression analyses examining associations between cardiovascular risk variables and corrected brain-age gap (winsorized at 2%) within the Gothenburg RA cohort (n=71). Each row represents a separate linear regression model including age and the indicated cardiovascular variable. Standardized β-estimates and 95% confidence intervals are shown. Upper panel: Framingham cardiovascular risk score and vascular age gap. Lower panel: individual Framingham score components, including smoking status, total cholesterol (Tot Cholesterol), high-density lipoprotein (HDL), systolic blood pressure (SBP), antihypertensive treatment (Hypertension drug), and diabetes status. **B)** Multivariable linear regression model of corrected brain-age gap including age, hypertension status, and antihypertensive treatment. Standardized β-estimates and 95% confidence intervals are shown. P-values are indicated for significant associations. **C)** The brain-age gap an RA patients stratified by hypertension status and healthy control cohorts. P-values was calculated with Kruskal-Wallis test with post-hoc comparisons. *p<0.05, **p<0.01, ***p<0.001, ****p<0.0001.

**Figure S4.**
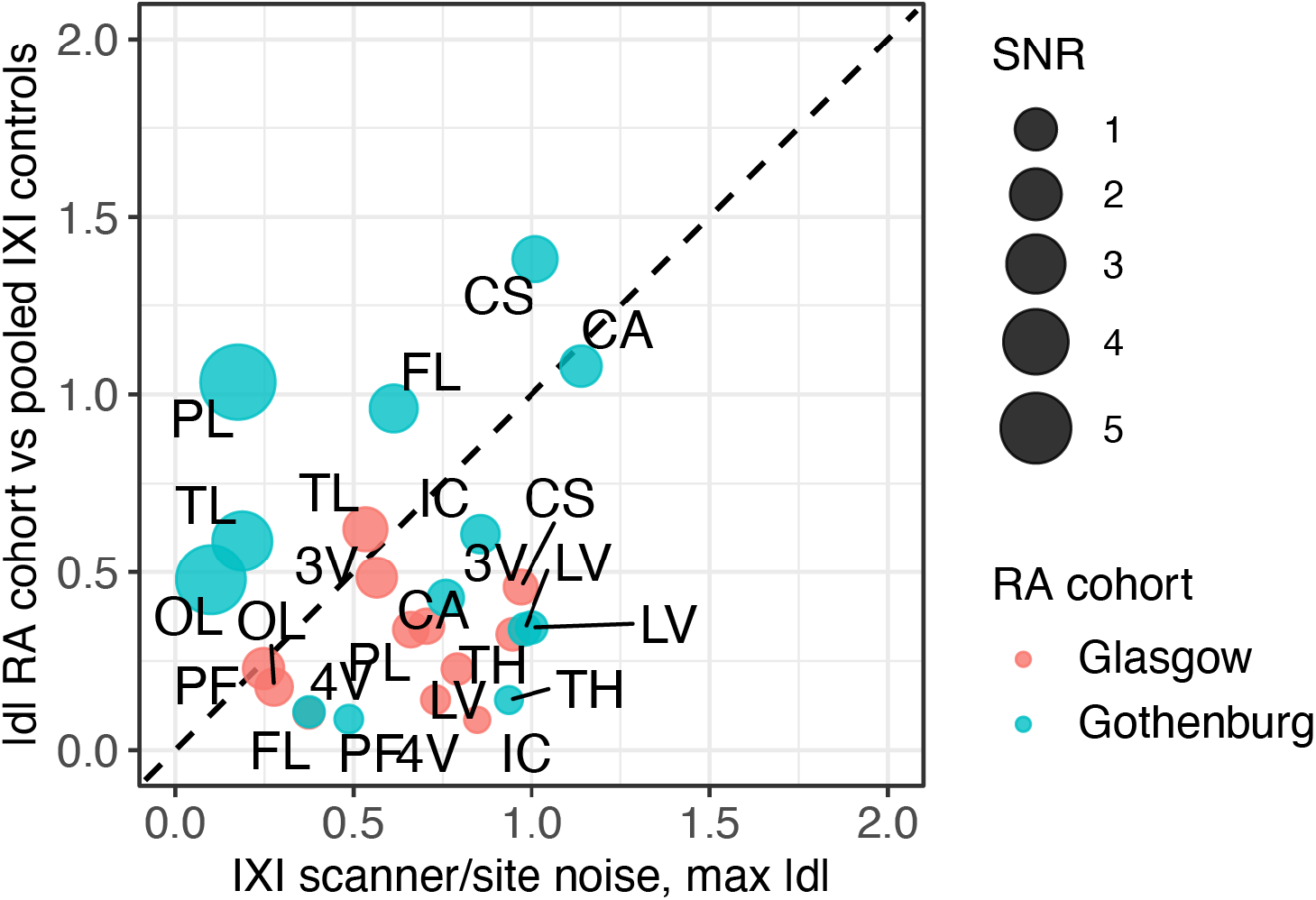
Regional brain volume differences in younger rheumatoid arthritis patients are limited relative to inter-site MRI variability. Brain region volumes derived from magnetic resonance imaging (MRI) were analysed in two independent RA cohorts (Gothenburg, n = 71; Glasgow, n = 50) and age- and sex-matched healthy controls without diagnosed neurological disease from three IXI sites (HH, Guys, and IOP). Controls were strictly matched for age (maximum difference 3 years). **A)** Scatter plot of absolute effect sizes (|d|, Hedges’ g) for differences between RA patients and pooled IXI controls under 60 years of age (y-axis) plotted against inter-site variability across IXI scanners (x-axis). Each point represents a brain region (abbreviations shown), colored by cohort and scaled by the signal-to-noise ratio (SNR = |d| / inter-site variability). The dashed diagonal line indicates equality between disease-related effects and scanner/site variability (|d| = noise). Points above the line represent regions where RA-associated differences exceed inter-site variability, whereas points below the line indicate effects smaller than scanner/site noise.

**Supplementary table 1.** Brain regions and super regions.

| Brain region name | Super region name |
| --- | --- |
| Medulla oblongata | Brainstem |
| Midbrain | Brainstem |
| Pons | Brainstem |
| Substantia nigra | Brainstem |
| Caudate nucleus | Central structures |
| Corpus callosum | Central structures |
| Mamillary body | Central structures |
| Nucleus accumbens | Central structures |
| Pallidum | Central structures |
| Putamen | Central structures |
| Subthalamic nucleus | Central structures |
| Thalamus | Central structures |
| Anterior orbital gyrus | Frontal lobe |
| Inferior frontal gyrus pars opercularis | Frontal lobe |
| Inferior frontal gyrus pars orbitalis | Frontal lobe |
| Inferior frontal gyrus pars triangularis | Frontal lobe |
| Lateral orbital gyrus | Frontal lobe |
| Medial orbital gyrus | Frontal lobe |
| Middle frontal gyrus | Frontal lobe |
| Posterior orbital gyrus | Frontal lobe |
| Pre-subgenual frontal cortex | Frontal lobe |
| Precentral gyrus | Frontal lobe |
| Straight gyrus | Frontal lobe |
| Subcallosal area | Frontal lobe |
| Subgenual frontal cortex | Frontal lobe |
| Superior frontal gyrus | Frontal lobe |
| Anterior cingulate gyrus | Insula & cingulate |
| Insula anterior long gyrus | Insula & cingulate |
| Insula anterior pole | Insula & cingulate |
| Insula anterior short gyrus | Insula & cingulate |
| Insula middle short gyrus | Insula & cingulate |
| Insula posterior long gyrus | Insula & cingulate |
| Insula posterior short gyrus | Insula & cingulate |
| Posterior cingulate gyrus | Insula & cingulate |
| Cuneus | Occipital lobe |
| Lateral remainder occipital lobe | Occipital lobe |
| Lingual gyrus | Occipital lobe |
| Angular gyrus | Parietal lobe |
| Postcentral gyrus | Parietal lobe |
| Superior parietal gyrus | Parietal lobe |
| Supramarginal gyrus | Parietal lobe |
| Cerebellum anterior lobe | Posterior fossa |
| Cerebellum corpus medullare | Posterior fossa |
| Cerebellum flocculonodular lobe | Posterior fossa |
| Cerebellum inferior posterior lobe | Posterior fossa |
| Cerebellum superior posterior lobe | Posterior fossa |
| Cerebellum vermis | Posterior fossa |
| Amygdala | Temporal lobe |
| Anterior temporal lobe lateral part | Temporal lobe |
| Anterior temporal lobe medial part | Temporal lobe |
| Fusiform gyrus | Temporal lobe |
| Hippocampus | Temporal lobe |
| Middle and inferior temporal gyrus | Temporal lobe |
| Parahippocampal and ambient gyri | Temporal lobe |
| Piriform cortex | Temporal lobe |
| Posterior temporal lobe | Temporal lobe |
| Superior temporal gyrus anterior part | Temporal lobe |
| Superior temporal gyrus middle part | Temporal lobe |
| Cerebral aqueduct | Ventricles |
| Fourth ventricle | Ventricles |
| Lateral ventricle | Ventricles |
| Third ventricle | Ventricles |

